# An Explainable Self-Attention Deep Neural Network for Detecting Mild Cognitive Impairment Using Multi-input Digital Drawing Tasks

**DOI:** 10.1101/2021.12.15.472738

**Authors:** Natthanan Ruengchaijatuporn, Itthi Chatnuntawech, Surat Teerapittayanon, Sira Sriswasdi, Sirawaj Itthipuripat, Thiparat Chotibut, Chaipat Chunharas

## Abstract

Mild cognitive impairment (MCI) is an early stage of age-inappropriate cognitive decline, which could develop into dementia – an untreatable neurodegenerative disorder. An early detection of MCI is a crucial step for timely prevention and intervention. To tackle this problem, recent studies have developed deep learning models to detect MCI and various types of dementia using data obtained from the classic clock-drawing test (CDT), a popular neuropsychological screening tool that can be easily and rapidly implemented for assessing cognitive impairments in an aging population. While these models succeed at distinguishing severe forms of dementia, it is still difficult to predict the early stage of the disease using the CDT data alone. Also, the state-of-the-art deep learning techniques still face the black-box challenges, making it questionable to implement them in the clinical setting. Here, we propose a novel deep learning modeling framework that incorporates data from multiple drawing tasks including the CDT, cube-copying, and trail-making tasks obtained from a digital platform. Using self-attention and soft-label methods, our model achieves much higher classification performance at detecting MCI compared to those of a well-established convolutional neural network model. Moreover, our model can highlight features of the MCI data that considerably deviate from those of the healthy aging population, offering accurate predictions for detecting MCI along with visual explanation that aids the interpretation of the deep learning model.

## Introduction

It has been estimated that, by 2020, approximately 50 million people will suffer from dementia worldwide.^1^ Unfortunately, there is currently no cure for such a devastating condition. One of the most crucial management strategies for preventing the disease from progressing is to detect the initial stage of pathological cognitive aging as early as possible. This initial stage of cognitive decline is known as mild cognitive impairment (MCI), which has a prevalence rate of about 20% of the elderly population aged 60 and above ^2–3^.

The clock-drawing test (CDT) is one of the most studied neuropsychological tasks known for its ability to capture a wide range of neurocognitive disorders including Alzheimer’s disease and other types of dementia. Accordingly, it has been included in a rapid population-based screening test for dementia^4^. While the CDT could be easily implemented in the pen-and-paper format, it still requires highly trained medical personnel to administer the screening as well as analyze and interpret the test results. Recently, research studies have tried to overcome these limitations by collecting the data in the digital format and adopting advanced machine learning (ML) models to automate and improve the scoring and disease classification methods ^5–15^. Initially, early ML research has demonstrated promising classification results obtained from domain knowledge feature construction guided by human experts ^5–7,10,14,15^. Moreover, recent studies have adopted deep learning models to alleviate the need for such hand-crafted features in several analytic steps including digit classification,^8^ digit-and-clock-hand recognition,^9^ contour-and-hand segmentation and digit classification,^13^ clock score prediction,^11^ and healthy-versus-cognitive-impaired classification.^12^ For example, compared to traditional ML techniques, deep learning techniques perform much better in classifying dementia versus health controls.

In contrast to the case of dementia, the success in using digital clock drawing and deep learning to detect less severe neurocognitive disorder like mild cognitive impairment is still limited. To improve the model performance, it is possible to combine multiple drawing tasks such as a trail making task or a copy drawing task as inputs to the deep learning model. However, the deep learning model is often referred to as a black box approach given that most of them provide only predictions without additional information that can be understood by humans. The model that hopes to be applied in the health-care system should be interpretable by health-case providers.^16^

In this work, we developed a novel multi-input deep learning model that integrates three different drawing tasks to perform an explainable MCI detection. Extending clock drawing-based detection with deep learning ^8,9,11–13^ to include a cube-copying drawing and a trail-making task into model inputs, our convolutional neural networks (CNNs) equipped with the self-attention mechanism (multi-input Conv-Att) achieve an excellent classification performance. The multi-input Conv-Att model enjoys an improvement of 0.051, 0.241, and 0.095 gain on the average accuracy, F1-score, and area under the receiver operating characteristic curve (AUC), respectively, compared to those of a baseline CNN. Importantly, with the self-attention mechanism, the model sensibly highlights features of each drawing tasks that deviate significantly from those of the healthy population, thereby providing multiple visual cues for medical practitioners to justify model predictions. While the prediction accuracy of the multi-input Conv-Att model is comparable to that of the baseline CNN with Grad-CAM^17^, the multi-input Conv-Att model provides visual cues that are more consistent with how clinical experts analyze drawing tasks. We also examine the standard medical criterion for MCI diagnosis - the drawings are classified as MCI with certainty if their score drops below a hard cutoff. Our data reveal that the scores of the healthy population and those of the MCI population strongly overlap, especially near the cutoff. Thus, we demand our multi-input Conv-Att model to output a class probability (i.e., soft labels), rather than a class with certainty, to accommodate the classification uncertainty near the cutoff. With soft labels, we further gain an improvement of 0.013 and 0.056 on the average accuracy and F1-score. Finally, in the spirit of reproducibility, we will make our dataset publicly available for interested researchers to benchmark their methods.

## Results

We assessed the MCI vs. healthy aging control classification performance of our proposed method, multi-input Conv-Att with soft label, on a dataset of 918 subjects (138 of which were used as unseen test data: 98 healthy aging controls and 40 MCI patients) acquired with informed consents at King Chulalongkorn Memorial Hospital, Bangkok, Thailand (see Material & Methods for more details). Each subject was categorized based on a Montreal Cognitive Assessment (MoCA) score cutoff of 25, ^18^ resulting in 651 healthy subjects and 267 MCI patients in the dataset. We compared the proposed method to a baseline model, denoted as single-input VGG16, that is closely related to existing deep-learning-based methods ^11,12^ on MCI vs. aging controls classification in terms of the framework used: input data, deep learning components, and a training procedure.

Since the proposed method strongly deviates from existing works by the incorporation of three specially designed components into a standard CNN, which consist of the multi-input approach (clock drawing, cube-copying and trail-making inputs), self-attention mechanism, and soft labeling technique, we also performed an ablation study to determine the relative improvement gained from each of the proposed components, as quantitatively measured by the accuracies, F1-scores, and AUC.

In addition to reporting the classification results in Table 1, we assessed the ability of the proposed method to support its classification decisions (MCI vs. healthy aging control) through visual interpretability. Particularly, we demonstrated that the proposed method yielded improved heat maps compared to those generated by the multi-input VGG16 model with Grad-CAM visualization,^17^ as measured by the interpretability scores given by 3 experts (a neurologist and licensed neuropsychologists) (Table 2).

**Table 1.** The mean and standard deviation of the accuracies, F1-scores, and AUCs over 5 different random training-validation-test data splitting. Our proposed model, which benefits from the incorporation of multiple complementary drawing tasks (clock drawing, cube-copying, and trail-making), self-attention mechanism and soft labeling approach, achieved much higher mean accuracy, F1-score, and AUC than the baseline single-input VGG16 model.

| Models | Accuracy | F1-score | AUC |
| --- | --- | --- | --- |
| VGG16 (with only CDT) | $0.7478 \pm 0.0071$ | $0.3573 \pm 0.0443$ | $0.7429 \pm 0.0131$ |
| Multi-input VGG16 | $0.7986 \pm 0.0071$ | $0.5938 \pm 0.0207$ | $0.8115 \pm 0.0192$ |
| Conv-Att (with only CDT) | $0.7522 \pm 0.0125$ | $0.3586 \pm 0.0309$ | $0.7337 \pm 0.0204$ |
| Multi-input Conv-Att | $0.7986 \pm 0.0071$ | $0.5981 \pm 0.0221$ | $0.8379 \pm 0.0176$ |
| Multi-input Conv-Att with soft label (proposed) | $0.8116 \pm 0.0103$ | $0.6539 \pm 0.0097$ | $0.8375 \pm 0.0116$ |

**Table 2.** The mean and standard deviation of the visual interpretability scores over all samples in the test set given by a neurologist and two licensed neuropsychologists (scores from 1 to 5; 1 being the worst and 5 being the best in terms of providing a visual interpretability that aligned with their experience and knowledge).

| Evaluators | VGG16 with Grad-CAM | Conv-Att with soft label (proposed) |
| --- | --- | --- |
| Expert 1 | 1.42 ± 0.74 | 3.41 ± 0.61 |
| Expert 2 | 1.86 ± 0.74 | 2.20 ± 0.93 |
| Expert 3 | 1.36 ± 0.62 | 2.87 ± 0.68 |

### MCI vs. healthy aging control classification

As shown in Table 1, the proposed method yielded the mean classification accuracy of 0.8116, F1-score of 0.6539, and AUC of 0.8375 over five repetitions, demonstrating 8.53%, 83%, and 12.7% relative improvements in the mean accuracy, F1-score, and AUC, respectively, with respect to the baseline method. By extending the single-input models (VGG16 and Conv-Att) to their corresponding multi-input models (multi-input VGG16 and multi-input Conv-Att), we observed significant improvement in the classification accuracies, AUCs, and, more remarkably, F1-scores. While the accuracies, F1-scores, and AUCs obtained from VGG16 and Conv-Att were comparable in both the single- and multi-input cases, the self-attention mechanism included in the Conv-Att model resulted in improved visual interpretability which will be discussed in the next subsection in detail. Including the soft label component further improved both accuracy and F1-score of the proposed model.

### Visual interpretability

An example of visual interpretability of the multi-input VGG16 model with Grad-CAM^17^ and the proposed model is shown in Figure 1. The quantitative evaluation of each method is presented in Table 2. The results clearly showed that the visual interpretability from our model is better than those obtained from the multi-input VGG model with the Grad-CAM method^17^.

**Figure 1.**
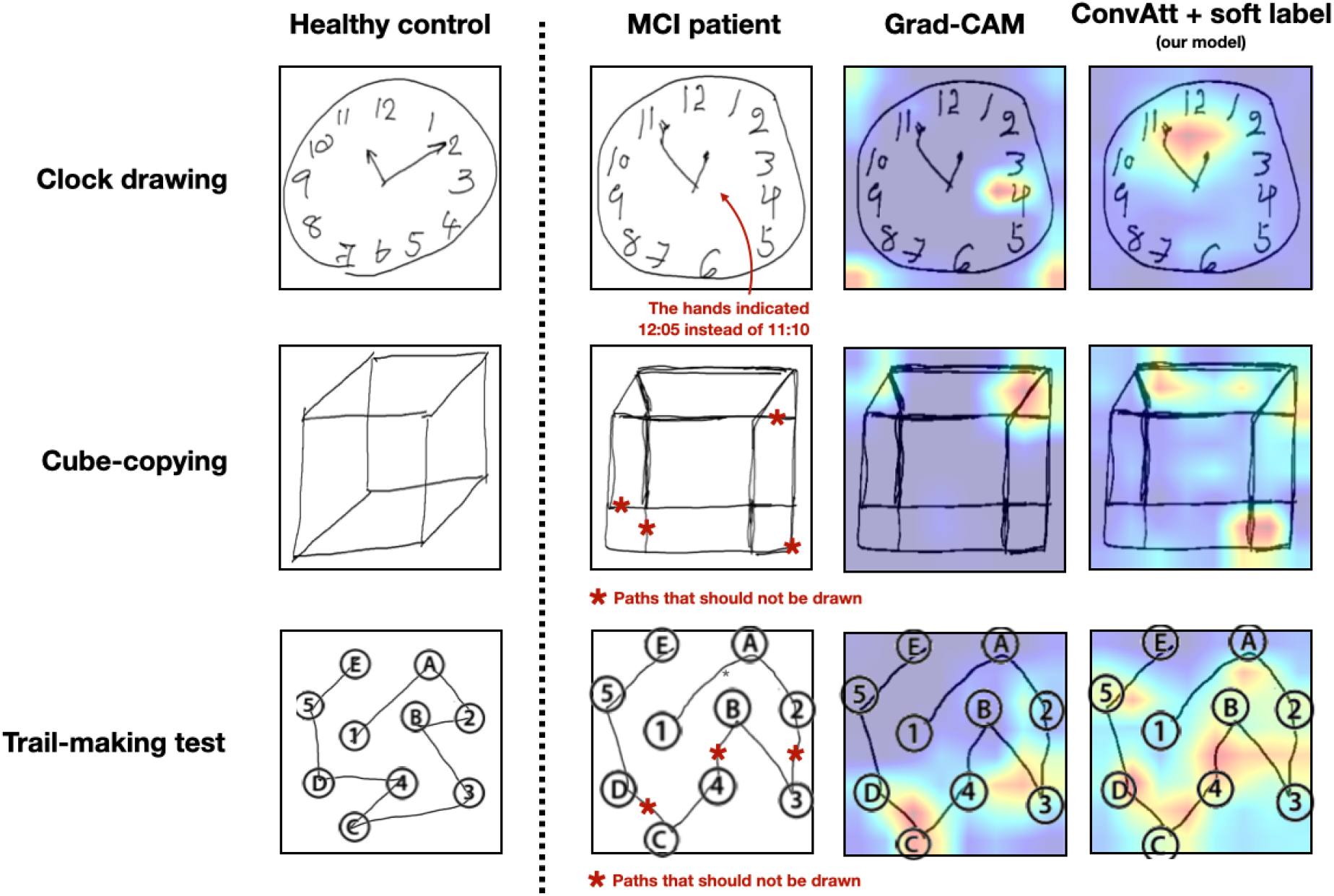
Visual explanations provided by the multi-input VGG16 model with Grad-CAM visualization (2^nd^ column from the right) and the proposed model (column on the far-right) on a representative MCI test sample (2nd column from the left). For the clock image (first row), our model highlights the hands of the clock where it says 12:55 instead of 11:10. For the cube-copying image (second row), our model highlights unnecessary paths better. For the trail-making task (last row), our model could focus along the paths that should not have been there (paths from 2-3, B-4 and C-D), while the multi-input VGG16 model with Grad-CAM failed to highlight some of those paths (B-4). Note that the red arrow and asterisks were not drawn by the subjects but added here to aid the descriptions.

## Materials and Methods

### Data collection

Under the institutional review board approval, a digital version of the MoCA test was administered on a tablet with a digital pen to a total of 918 subjects with informed consents by trained psychologists at King Chulalongkorn Memorial Hospital, Bangkok, Thailand. The population came from a healthy elderly cohort which focuses on preventive care for healthy Thai citizens without major medical conditions (such as organ failures). The median age was 67 years old (ranging from 55-89 years old), 77% female, 44% received bachelor’s degree, and 20% received higher education. For the clock drawing task, the subjects were instructed to “draw a circular clock face with all the numbers and clock hands indicating the time of 10 minutes past 11 o’clock”. In the cube-copying task, the subjects were instructed to copy the Necker cube image on an empty space. In the trail-making test, the subjects were instructed to “draw a line that goes from a number to a letter in an ascending order, starting at number 1 (pointing to the number 1), to this letter (pointing to the letter A), then go to the next number (pointing to the number 2), and so on.”

From each subject, we extracted the drawn clock drawing, cube-copying and trail-making images along with the MoCA score. We then categorized the subjects into healthy aging controls and MCI patients based on the MoCA score. In particular, the subjects were categorized as having MCI if the MoCA scores were below 25 as typically used in clinical routines^18^, resulting in 651 healthy subjects and 267 MCI patients in our dataset. The collected data were randomly split into three groups in a stratified fashion: 70% as training, 15% as validation, and 15% as test data. All images were resized to 256×256 in all experiments.

### Proposed method: multi-input Conv-Att model with soft label

We developed a multi-input deep learning method for MCI vs healthy aging controls classification that is a cascade of convolutional neural network (CNN) backbones and self-attention layers^19,20^ trained with soft labels, as shown in Figure 2. As opposed to existing models which take a clock drawing image as the only input to the models,^11,12^ our proposed multi-input model takes clock drawing, cube-copying and trail-making images simultaneously as inputs, exploiting complementary information offered by the three neuropsychological tasks. Incorporating the self-attention layers into the model leads to more efficient image representations, compared to typical CNNs, that can later be used to support the model’s classification decision through heat-map visualization. The soft-label component of our method is designed to aid our model training by taking into account the uncertainty of the diagnostic labels (i.e., MCI vs. healthy aging control) near the designated MoCA score cutoff. In the following subsections, we described each of the components in detail.

**Figure 2.**
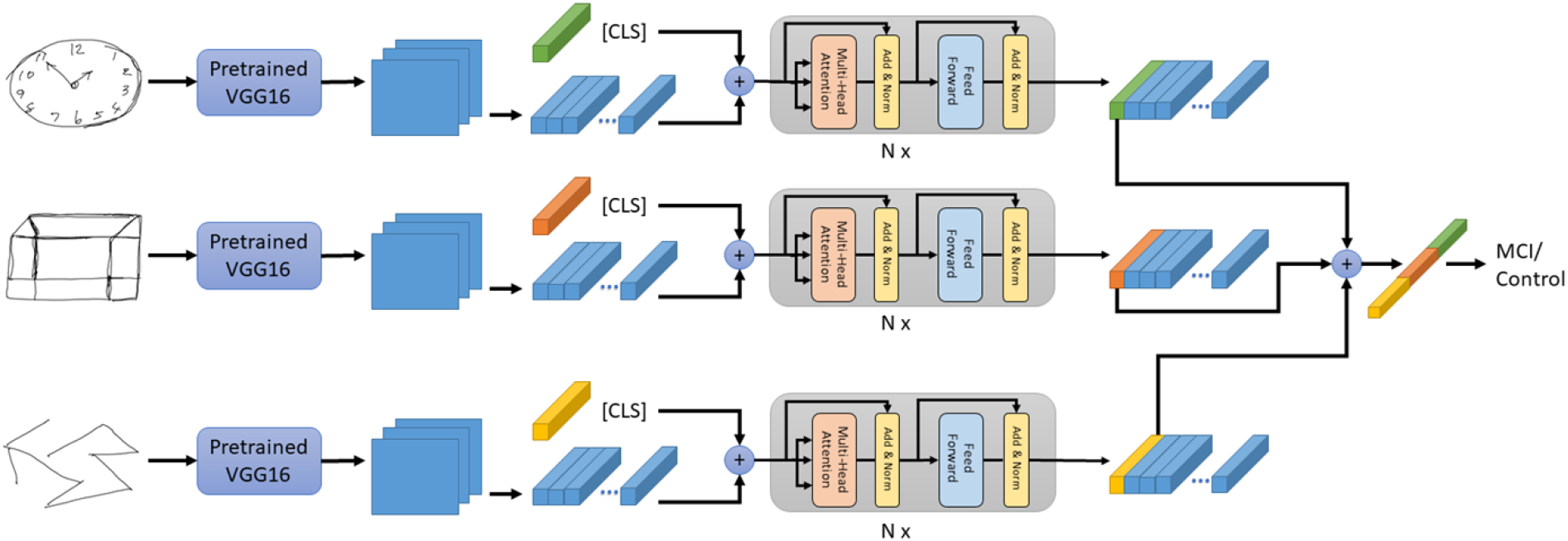
Overview of our proposed multi-input Conv-Att model. Our model simultaneously takes clock drawing, cube-copying, and trail-making images as its inputs and processes them using a cascade of CNNs and a stack of self-attention layers.

#### Conv-Att model architecture

As shown in Figure 2, clock drawing, cube-copying, and trail-making images are used as the inputs to our model. Each of the three images is passed into a separate CNN backbone (VGG16 ^21^ pretrained on the ImageNet dataset,^22^ followed by a stack of self-attention layers, resulting in a vectorized image representation. Including the self-attention layers to the model leads to not only efficient image representation, but also improved visual explanation for MCI vs. healthy aging controls classification. Then, the resulting vectors from the three tasks are concatenated and processed by a two-node fully connected layer with the softmax function

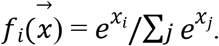

#### Self-attention

Unlike in a standard image classification model, we employ self-attention instead of a pooling layer to aggregate the output from the CNN backbone, *X* ∈ *R*^*H×L×C*^ where *H,L,C* are the height, width and number of filters. First, we initialize a random classification token vector, [CLS] ∈ *R*^*D*^ where *D* is the hidden dimension in the self-attention mechanism used in BERT.^23^ The [CLS] vector is used to aggregate visual representation from all pixels in *X*. Second, 1×1 convolution with *D* output filters are applied to *X* to adjust its last dimension to match the hidden dimension *D* of the [CLS] vector. After that, it is reshaped into a matrix of shape HL × D and concatenated with [CLS] resulting in 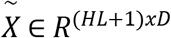. Reshaping the data this way enables us to investigate how much each pixel contributes to the final classification decision through the attention rollout method.^24^ The self-attention is defined as:

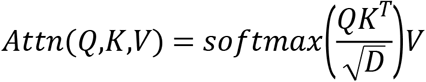

where *Q,K,V* ∈ *R*^(*HL*+1)×*D*^ are the query, key and value which are the projections of 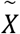 with different linear function: 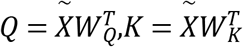, and 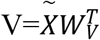 where *W*_*Q*_, *W*_*K*_, *W*_*V*_ ∈ *R*^(*HL*+1)×*D*^. At the final layer of self-attention, the vector at the [CLS] position is used as the final image representation.

#### Soft-label

As explained in the Data collection subsection, we assigned the label of 0 (healthy control) to a subject with the MoCA score higher than or equal to 25, and the label of 1 (MCI) otherwise. Such a labeling approach is typically referred to as hard labeling. While training a model with hard labels is the most commonly used approach to solving binary classification, we propose to train our proposed model with soft labels based on MoCA scores for MCI vs. healthy aging controls classification to take into account the uncertainty of the diagnostic labels (MCI vs. healthy aging control) near the MoCA score cutoff of 25. Specifically, we define a soft label *y* according to the following equation

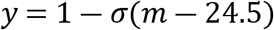

where *m* is a MoCA score, and *σ* denotes the sigmoid function. Since a subject with the MoCA score of 24 is labeled as an MCI patient and a subject with the MoCA score of 25 is labeled as a healthy control, we subtract 24.5 from *m* so that the center of the sigmoid will be at 24.5.

The hard threshold of 25, below which a subject is considered an MCI patient, is a man-made criterion, rather than the number revealed through rigorous statistical tests from a large number of trials and can be varied depending on contexts such as education or cultures.^18,25,26^ Rather, by assigning a soft label, the uncertainty in the classification result is manifested through the sigmoidal probability function. So, in a post hoc way, the soft label approach can help relax the strong classification bias inherent in the hard label approach.

We trained the proposed model by minimizing the binary cross-entropy loss

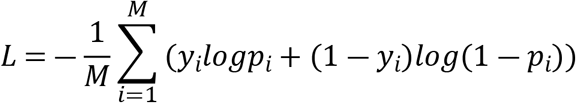

where *M* is the number of training data, 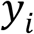is the soft label of the data 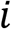, and 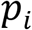is the output of the model which can be interpreted as the predicted probability that the data 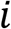 is associated with MCI.

#### Attention rollout

To visualize how self-attention combines the pixels of the last feature maps calculated by the CNN backbone *X* into the final image representation, we used attention rollout^24^. In the self-attention layers, there exist residual connections between consecutive layers. Therefore, the output of the self-attention layer *l* +1 is defined below:

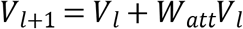

where *W*_*att*_ is the attention weight, and *V*_*l*_ is the output of the self-attention layer *l*. To compensate the effects of the residual connections, the raw attention *A* is *A* = 0.5*W*_*att*_ +0.5*I* where *I* is the identity matrix. The attentions from self-attention layer *l*_*i*_ to layer *l*_*j*_ are computed by recursively multiplying the attention weights as follows:

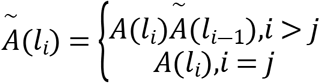

where 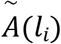 is the attention rollout at self-attention layer *l*_*i*_, and *A*(*l*_*i*_) is the raw attention at self-attention layer *l*_*i*_. The interpretability from our model is how the self-attention layers combine the last feature maps into the final image representation. Therefore, it is equivalent to the attention rollout for [CLS] over all the pixels of the last feature map. The attention rollout for each [CLS] is reshaped back to the size of the last feature map and then is resized to match the size of the original input image. The heat map from each [CLS] is used as the interpretability for each input image.

## Experiments

We compared our proposed method to two VGG16-based models: a single-input VGG16 model that takes only a clock drawing image as input and a multi-input VGG16 model that simultaneously takes clock drawing, cube-copying and trail-making images as inputs. For the multi-input version, different VGG16s were used to process different input images. At the end of each VGG16, the global average-pooling layer was applied. The average-pooled image features from each task were concatenated and then passed into a two-node fully connected layer with the softmax function.

We also compared the results of the proposed method to those of the proposed method with some components removed. In particular, we recorded the performances of (1) the single-input Conv-Att model and (2) the multi-input Conv-Att model, both trained with hard labels.

### Model training

Adam^27^ optimizer with the learning rate of 1e-5, *β*_1_ = 0.9,*β*_2_ = 0.99 and *ϵ* = 10^−7^ were used in all experiments. The models were trained for 100 epochs with the batch size of 64. We adopted image augmentation to increase the effective size of the training data. Specifically, we first zero-padded the image to size of 280×280, and then cropped the image back to 256×256 with the center at a random location in the padded image. For the models that included stacked self-attention layers, we used 3 self-attention layers with the number of heads of one, hidden dimension size of 128, and hidden dimension size of the feedforward layer of 512.

### Evaluation

We performed 5 random training-validation-test data splitting and reported the mean and standard deviation of the accuracies and F1-scores obtained from each method. Since all the methods were trained to predict the probability of having MCI, *p*, for each input, we needed to convert the model prediction into a diagnostic label (i.e., hard label) so that we could compute accuracy and F1-score meaningfully. In this case, we categorized all the subjects with *p* ≥ 0.5as MCI patient and *p* < 0.5as healthy control. We also reported the AUCs for all the models under consideration.

We also assessed the ability of the proposed method to provide visual explanation to support its diagnostic decision by comparing the heat maps generated by the proposed model to those generated by the multi-input VGG16 model with Grad-CAM,^17^ which is one of the most commonly used methods for visual explanation. For each subject, we obtained the heat maps from both the proposed method and the multi-input VGG16 model. Then, we displayed them side-by-side and separately asked 1 neurologist and 2 licensed neuropsychologists to give scores between 1 and 5 to each set of heat maps (1 being the worst and 5 being the best in terms of providing visual explanation that aligned with their experience and knowledge). To avoid potential bias, we randomly shuffled the display locations (left vs. right) in a way that the heat maps of the proposed method were displayed on the left and the right of the VGG16 model with equal probability.

## Discussion

Incorporating three specially designed components into a standard CNN, consisting of the multi-input approach, self-attention mechanism, and soft labeling technique, resulted in our proposed model that outperformed the baseline CNN model as measured by not only several quantitative evaluation metrics (accuracy, F1-score, and AUC), but also the visual explanation provided by the model’s heat maps. Unlike in the previous studies that take only a clock drawing image as the input to the model,^11,12^ our proposed multi-input model exploits the complementary information provided by the three drawing tasks that rely on different combinations of fundamental cognitive abilities under the neuropsychological perspective: planning and task-switching in the trail-making test, visuospatial ability in the cube-copying drawing test, and both planning and visuospatial ability in the clock drawing test. By replacing the global max-pooling layer of a standard CNN architecture with a stack of self-attention layers, the proposed model was able to represent the data more efficiently and provide improved heat maps that could be used to support the model’s classification decision, compared to the baseline model visualized using Grad-CAM^17^. In addition to the multi-input and self-attention components, the proposed model benefits from the soft labeling technique that takes into account the uncertainty of the diagnostic labels (i.e., MCI vs. healthy aging control) near the MoCA score cutoff, resulting in further improvement in the classification performance.

While typical model evaluation tends to focus on classification performance, there are some other important aspects of what counts as a good model. In our case, we found that even though applying self-attention did not lead to better classification performance, clinical experts preferred the visual explanation provided by our model to those of multi-input VGG16. The clinical experts found that our model was able to highlight the areas that aligned with their experience and knowledge. For example, our model was able to highlight the clock hands where the locations of the short and long hands were incorrectly drawn (see Figure 1). This improved interpretability will become even more critical when AI is used as a decision support system - the direction that is gaining more attention these days.

In this study, we benchmarked our proposed model to not only a strong VGG16 baseline, but also several extensions of the baseline through our ablation study. Nevertheless, it is not straightforward to compare our reported quantitative metrics to those reported in existing works due to many reasons. First, different proxies of neurocognitive disorders have been adopted in different studies. For example, while the Shulman clock scoring system^28^ has been used as a surrogate marker for dementia (Chen et al., 2020), we used Petersen criteria with the MoCA score cutoff of 25 ^2,18^ for the MCI diagnosis in our study. Moreover, different study populations with potentially different demographic information and recruiting procedures were involved. Such experimental inconsistencies make it difficult for direct comparisons between the studies (for example, our data came from our healthy geriatric clinic where the MCI patients have only mild symptoms), warranting the sharing of the datasets of each study for better benchmarking. Consequently, we will make our dataset publicly available.

Switching the MoCA test from pen-and-paper to a digital format allows us to store the drawing trajectory of each drawing task, which contains both temporal and spatial information. While the proposed model, which only uses the final drawing as its input (only the spatial information) and discards the information-rich temporal information provided by the drawing trajectory, already achieved much higher accuracy, F1-score, and AUC than the baseline model, we project that its extension to a spatio-temporal version would further improve the classification performance with the presence of a larger amount of data. Since there exist many drawing trajectories that correspond to the same final drawing, having access to the raw drawing trajectory would enable the model to come up with potentially better data representation in the spatio-temporal domain. Moreover, exploring hidden structures in the high-dimensional space of raw drawing trajectory using unsupervised learning is also an interesting future direction.

In summary, we found that, in a challenging scenario where the aim is to identify mild MCI patients among healthy aging, using multiple inputs to train the model with soft labels and the self-attention mechanism leads to substantial improvements in both model performance and interpretability - both of which are important aspects of AI in medicine.

## Acknowledgements

The study received funding #SUNN64002 from Foundation of Thai Gerontology Research and Development institute (TGRI) for data collection and from National Research Council of Thailand for data analysis.

